# Characterization of an ant colony (Paraponera clavata) during rest and relocation phases: an experimentally induced protocol

**DOI:** 10.1101/2022.07.03.498617

**Authors:** Alexandra Rodriguez, Dominique Fresneau, Guénaël Cabanesa

## Abstract

Understanding societies of individuals is a challenging task. In several ant species individuals seem to have the same physical characteristics and, in principle, could assume any role the social environment requires. However, most ants’ societies present a caste organization in their colonies with particular roles. For the biologist understanding the complex dynamics ruling a colony is hard due to the difficulty of collecting and classifying long term ant activities in the field. The relocation phenomena which generally correspond to perturbation phases suffered by ant colonies are critical moments in the colony history and require rapid and effective response from workers in order to ensure the survival of the colony and its setup in a new safe nest. How the colonial group deals with this process? Is there a particular organization of the workers during the relocation periods? Here, we applied an experimentally induced protocol in the laboratory combined with behavioral observations and hierarchical classifications in order to identify social roles in a ponerine species, *Paraponera clavata*, and the repartition of labor during two resting periods and two relocation phases. The method provided significant information concerning the social dynamics of ant colonies and contribute to understand how species copes with a novel and stressing situation conducing to the occupation of a novel available nest.

## Introduction

In animal groups, parental behaviors are considered as survival and reproductive strategies, which increase fitness. In order to optimize their reproductive investment, several animal species live in groups. Social life implies cooperation between individuals. Interactions between individuals are based on several communication signals which allow them to assume particular roles in the social organization (Möglich 1978). This is the case of ants’ societies where all species have reached high levels of sociality, eusociality, and are characterized by three fundamental aspects: cooperation in young care, at least two generations overlapping and division of labor (Wilson 1971). This last characteristic is known as polyethism and it is based on the existence of ethological castes specialized in the accomplishment of particular tasks (Oster and Wilson 1978).

In ants, polyethism can be related to size polymorphism or to workers’ age. It can also depends on the society size (Lenoir 1979). Thomas and Elgar (2003) have observed that in small colonies young and old workers shared equally the time invested in activities conducted inside and outside the nest. On the contrary, in big colonies, old workers are specialized in food storage whereas young ones take care of the brood.

Ant colonies may be nomads or sedentary. However, all the sedentary species even those investing in big nests are able to move to novel nesting sites if they suffer important environmental perturbations. Behavioral mechanisms that take place during the emigration are not very different from those of nomad species (Porter and Bowers 1980; Wilson 1980; Fowler 1981; Hölldobler and Wilson 1990). In fact, all the species that have been studied for their nest stability have been shown to be able to move to novel areas independently of the nesting sites complexity. For instance, in species nesting in ground twigs, like *Myrmica punctiventris* or *Leptothorax longispinosus*, three quarters or almost all the nests may be abandoned during the year (Herbers 1985, 1986). In natural conditions (excluding budding and colonial fission observed in species that have not queens) five factors have been listed that can enhance a colony to move to a novel nesting site:

1. Nest perturbation: A mechanical perturbation or an inundation induce an immediate move of the queen, of the young workers and of the brood towards the intact chambers whereas foraging workers explore actively the neighborhood in order to found a safety and protected site where the colony can migrate. *Leptothorax colonies* react rapidly to any light presence in the nest, particularly the brood chambers, and transport all the colony members to a darker novel site (Möglich 1978).
2. Changes in microclimatic conditions: Even if there are not major perturbations, a progressive degradation of temperature and humidity conditions in the colony can induce a definitive nest abandon. This is the case of *Pachycondyla apicalis* and *Pachycondyla verenae* two ponerine ants that live in ephemeral nests formed by dead wood in the litter. Pezon and collaborators (2005) observed that queens were systematically recruited in the middle of colonial migrations optimizing their protection. Artificial shade or shade resulting from surrounding vegetation growth can be enough to induce moving in colonies of *Iridomyrmex purpureus* (Greenslade 1975 a,b), *Pogonomyrmex badius* (Carlson and Gentry 1973) and *Aphaenogaster rudis* (Smallwood 1982).
3. Predation and competition: Nest invasion by a predator or repeated attacks by a competitor induce nest abandon, sometimes in a catastrophic way (Wilson and Regnier 1971). When submitted to an invasion of legionary Neivamyrmex ants, individuals of *Pheidole desertorum* colonies evacuate totally their nest (Droual and Topoff 1981, Droual 1983). Typically, colony’s members disperse in all directions as a panic response and regroup in unoccupied sites (Droual 1984). When submitted to homospecific or heterospecific pression, harvester ants from Pogonomyrmex genus emigrate and establish a new nest in less occupied zones (Hölldobler 1976; De Vita 1979; Harrison and Gentry 1981; Gordon 1992). Emigration may be due to a resources and prey decline in the colony foraging area.
4. Seasonal polycalism: Species like *Polyrhachis dives* (Yamauchi et al. 1987), *Leptothorax ambiguous* (Herbers and Grieco 1994) and *Cataglyphis iberica* (Cerdá et al. 2002) expand their foraging area by dispersing to several nesting sites. Contact between sites is maintained by exchanges between foraging members and by brood transport. These particular emigrations save social colony cohesion and allow an effective resources distribution.
5. Superiority of a new site: Colonies may migrate in the absence of an imminent reproduction period and before any deterioration or perturbation of the nest. This migration may allow the appropriation of a better available nesting site. This kind of strategy has been recently observed in *Leptothorax albipennis* (Dornhaus et al. 2004) and significantly reduces the inherent emigration risks. It allows foraging ants to select the best site without time pressure and maintaining the colony protected in the ancient nest.

Even if colonies emigrations are widespread and frequent phenomena it stills be a dangerous activity for the whole colony. By consequence, all strategies minimizing the queen and the brood exposition is essential to the whole colony survival (Pezon et al. 2005).

The main objective of this study was to understand the logistic organization of the migration in *Paraponera clavata* and in particular to detect which individuals conduct the brood transfer from the old to the new nest.

*Paraponera clavata* is a good model for this research project as it is one of the rare ponerines which is both monogyne and polydomic as Hypoponera bondroiti, *Pheidole berthouldi, Pheidole goёldi a*nd *Pheidole hottentota* (Debout et al. 2007).

The collective and individual decision rules followed by workers as well as the specific mechanisms underlying colony organization in this species are still poorly understood so it is essential to find the rules that govern ant individual behavior and its integration at the colony scale.

We choose to apply an RFID method (Cabanès and Benani 2010) which allowed us to follow the position of each ant of the colony in the experimental dispositive which was composed by a nest area with several chambers, a foraging area, and a potential new nest site.

We monitored individual activities using passive radio-frequency identification (RFID) technology, a novel procedure applied to ants. RFID tagging has no observable effects on the behavior of the ants (Robinson et al. 2009, Cabanès and Benani 2010) and allows tracking the activity patterns of large numbers of individuals over a long period. The results of this automatic monitoring are presented in another publication. Here we analyse the results obtained by direct observation of resting periods and artificially induced relocations.

The fundamental issue investigated here is to understand how individuals can undertake the activities (nursing, transporting, foraging) according to the emerging needs of the colony. We wonder how these activities are distributed in the colony and if there are social castes or labor roles in a ponerine species in which anatomical distinction of individuals is very difficult because they do no present morphological dimorphism facilitating the recognition of clear castes.

## Material and methods

### Biological material

#### The model species

We followed and analyzed the movement of a ponerine ant colony composed of great sized ants easy to mark and follow. Ponerine ants are tropical species whose colonies consist in a few hundred to a few thousand individuals (Peeters, 1993). Paraponera clavata, whose distribution corresponds to the northern portion of South America (Bolton et al. 2005), nests under the trees and specially at the base of Pentaclethra macroloba that founding queens search actively (Bennet and Breed 1985, Hölldobler and Wilson 1990). The worker caste is lightly polymorphic with the onset of allometry and this polymorphism reflects a polyethism where the small workers are involved mostly in brood care, while the large ones are guardians and foragers (Breed and Harrison 1988, Breed 2002). Paraponera clavate feeds on nectar and small arthropods that it hunts actively foraging mainly in the canopy (Young and Hermann 1980). The species uses chemical recruitment to food sources applying individually discriminable pheromone trails (Barret et al. 1985, Breed et al. 1987, Breed and Harrison 1987).

In these species, queens are often not very different from workers, but a wide variety of social structure exists, ranging from monogyny to polygyny, with some species even lacking a specific queen caste (Peeters, 1997; Monnin and Peeters, 1998). Our studied species, *Paraponera clavata*, establishes monogyne colonies of up to 2500 monomorphic workers (Hölldobler, 1984).

#### Collection and housing of the studied colony

Th colony that we studied was collected in Belèm (Brazil) in December 2008. It included a queen, 3 wingless females and about 200 workers (due to the death occurred during transportation and installation in the laboratory the colony contained 110 workers in February 2009. A serial of observations were conducted in 2009 and 2010 that will be presented in another publication.

The study we presented her was conducted in 2011 when the colony was composed of 95 workers, one queen and one male.

The colony was placed in a nest composed of tree plastic boxes (dimensions: 77/115/13 mm each) with a floor of thick plaster of Paris. The boxes were connected by PVC pipes and covered with red gelatine covers ensuring the darkness for ants (since ants cannot see the red part of the spectre), while allowing observations (Figure 2 and 3). The nest was connected to the arena (69/48 cm) whose edges were coated with Fluon to prevent escape. The arena was connected to another nest composed of tree plastic boxes closed and unused by the ants. The floor of the arena was covered with a thin layer of sand. The ants were fed three times a week (Monday, Wednesday and Friday) with meal worms and apple-honey ad libitum. Water was present all the time on the arena in small dishes with absorbent paper. The temperature in the laboratory was maintained around 25°C.

**Figure 1:**
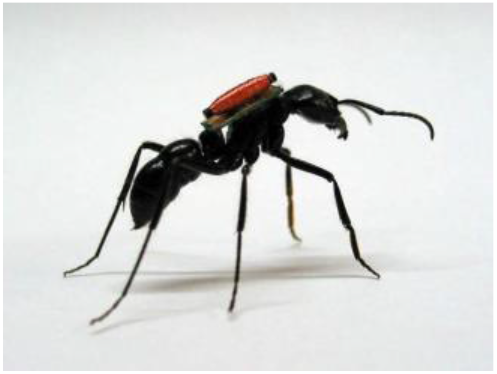
Ant marked with an RFID tag

**Figure 2:**
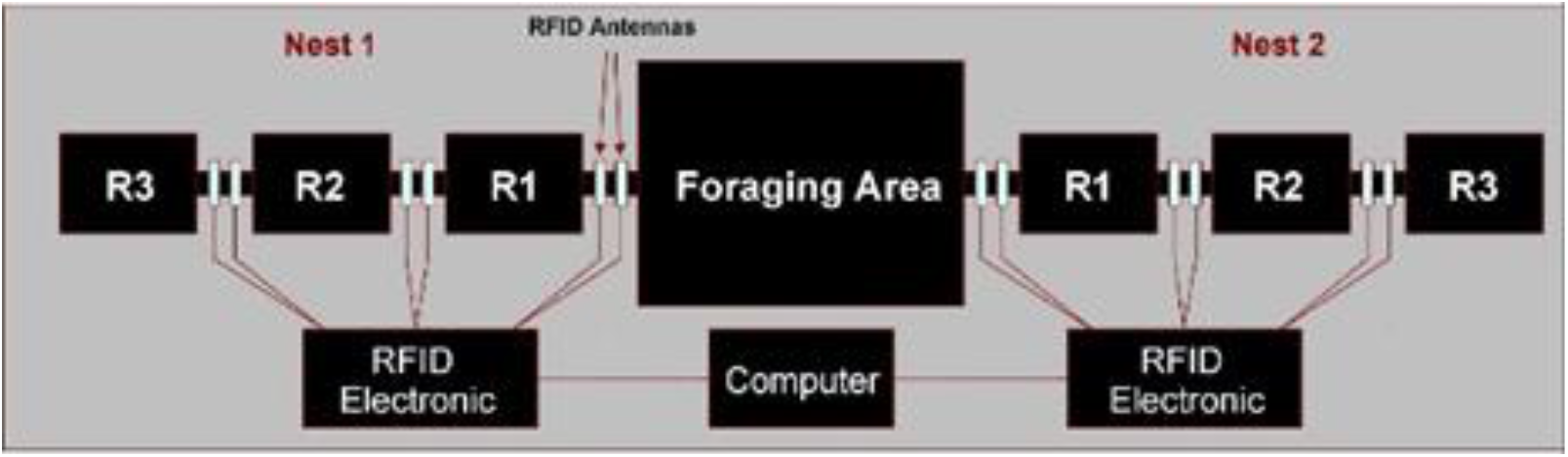
General view of the experimental device

**Figure 3:**
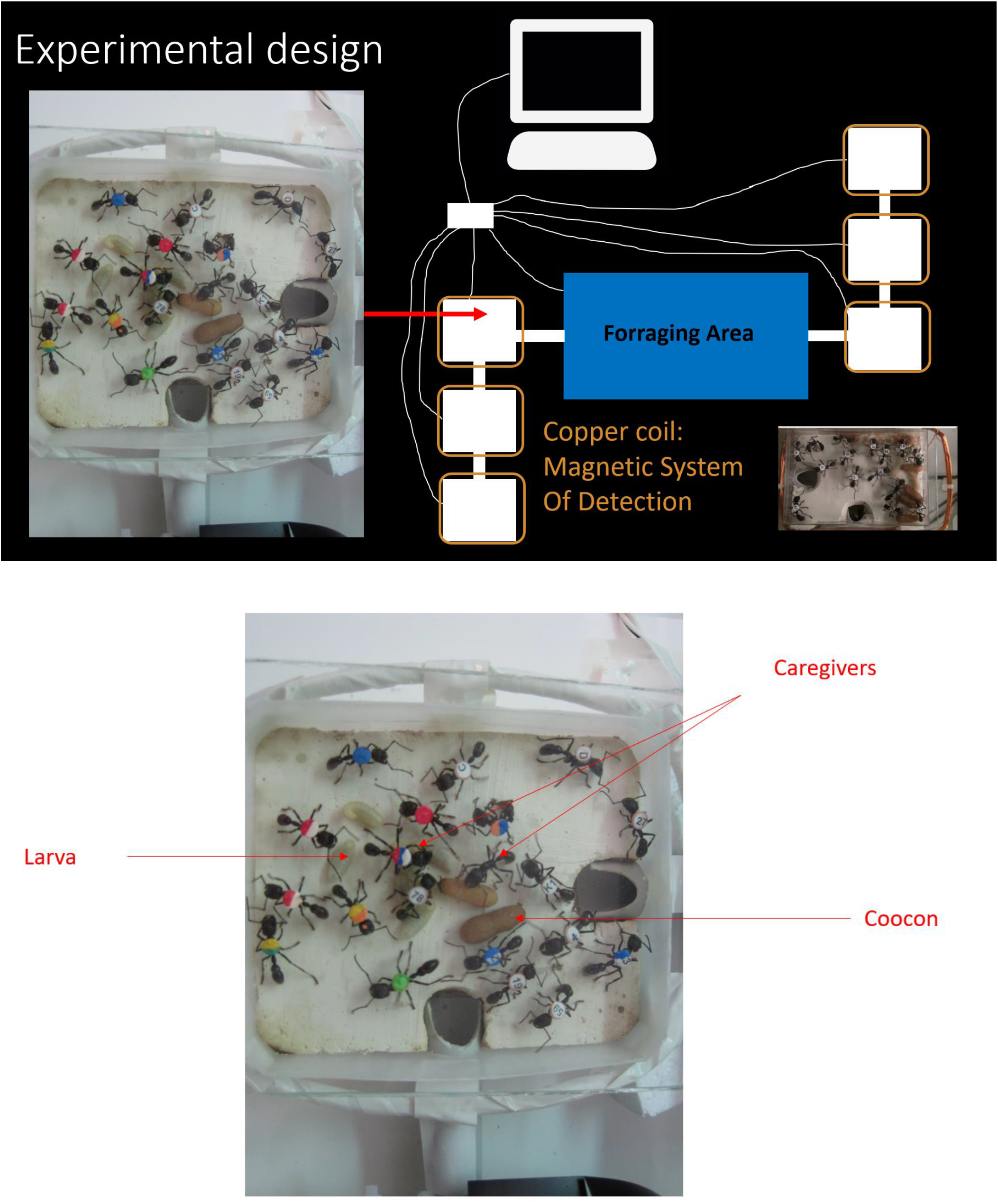
Detailed views of the experimental device

**Figure 4:**
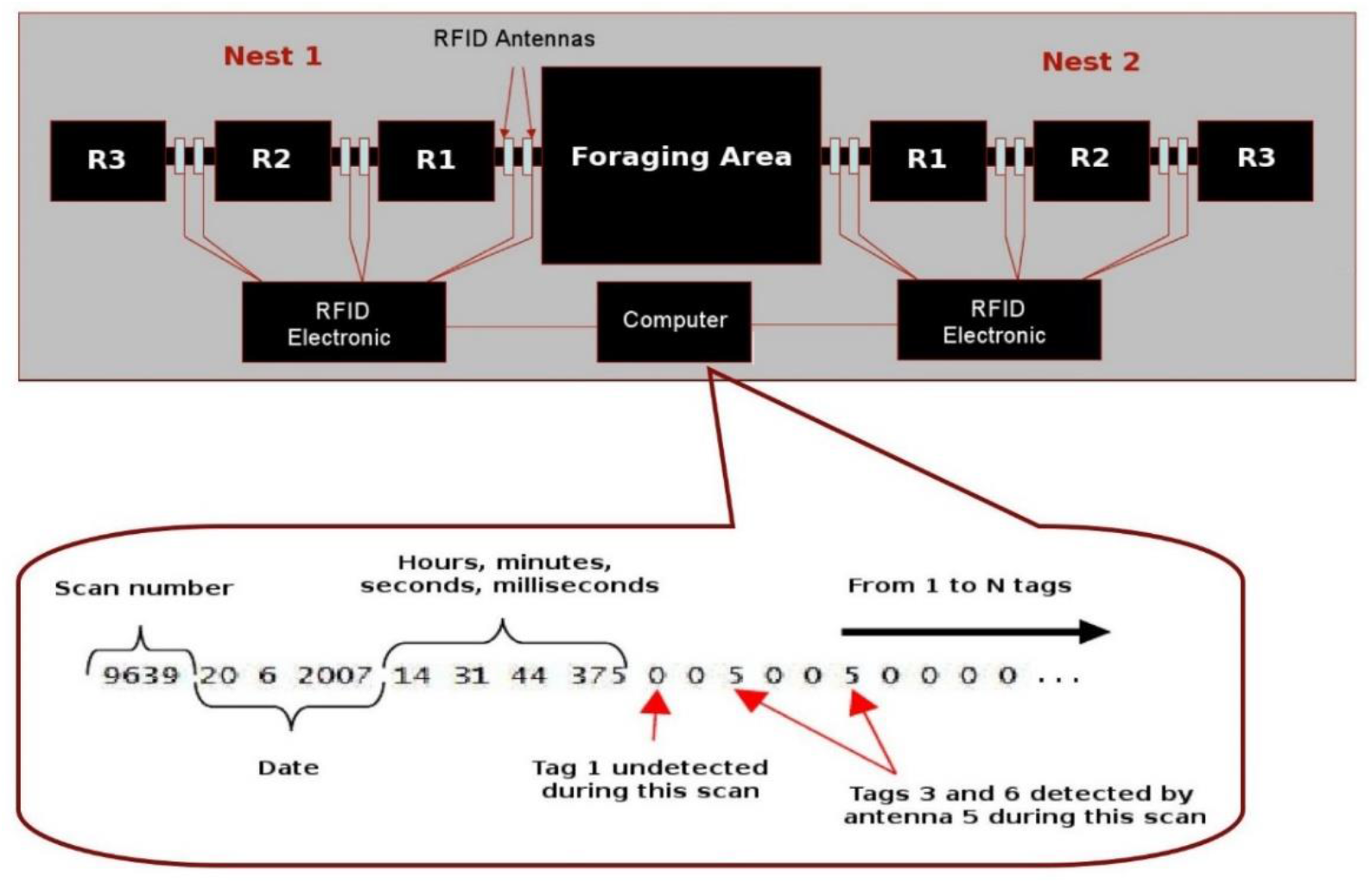
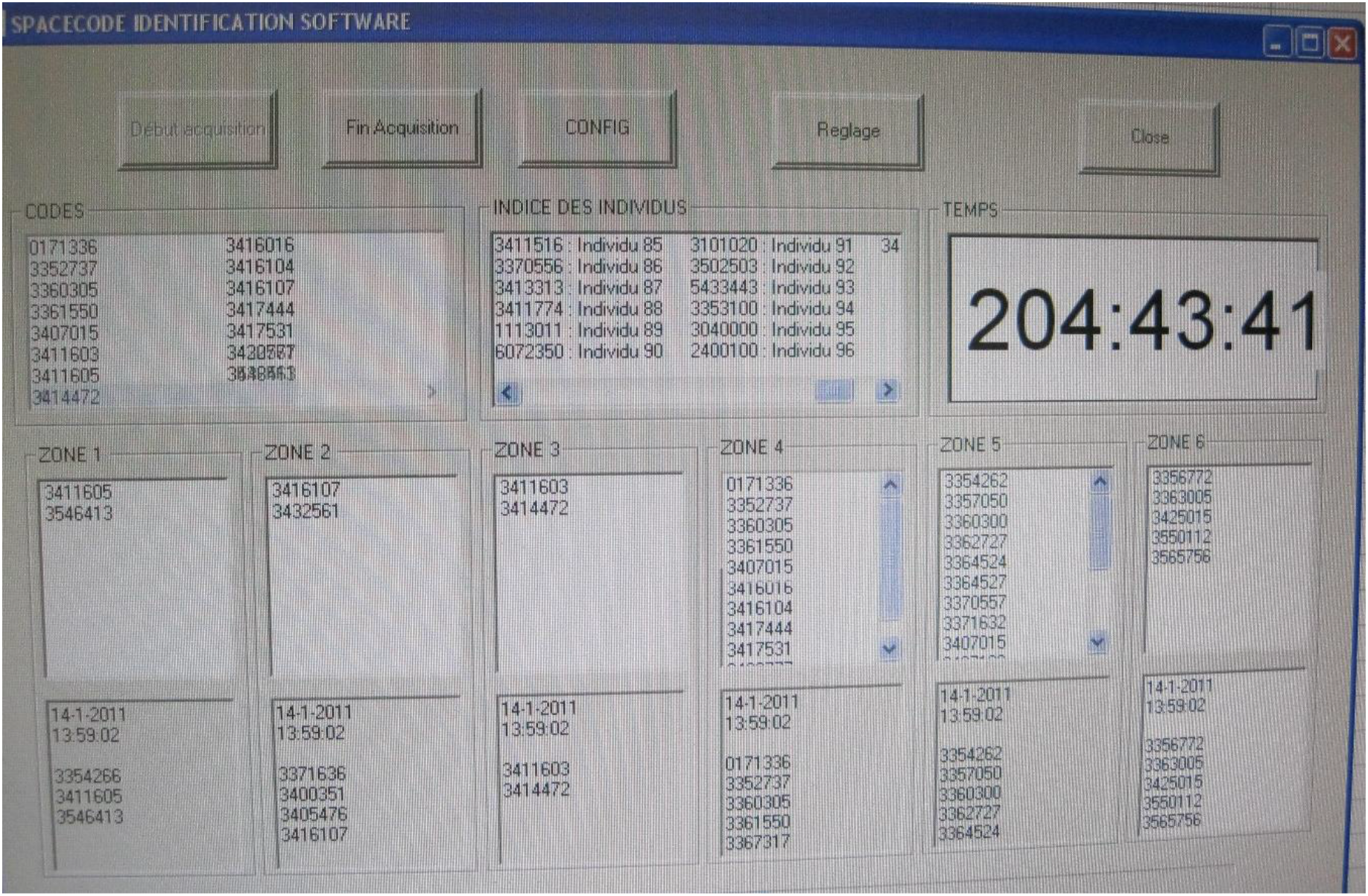
Example of a recorded scan in the data file

#### Ant marking and RFID-detection set-up

All adult ants except the queen were marked with labels composed of an RFID chip with plastic pellet with a number or color code allowing visual identification. Preliminary tests showed that the tags don’t disturb the ant behavior and the colony dynamic significantly.

The chip was coated with cyanoacrylate glue assuring mechanical protection. The labels were attached to the dorsal face of ants’ thorax using cyanoacrylate glue. Each chamber and the exit of the nest were equipped with antennas operating at 124 Hz frequency, allowing RFID detection continuously (every 10 s approximately). RFID chips and antennas have been provided by SpaceCode (www.spacecode-rfid.com)

Videos of the workers activity may be consulted on youtube with the following links:

Paraponera clavate taking care of the brood:
https://www.youtube.com/watch?v=hCK2QZn0QHU
https://www.youtube.com/watch?v=eJudnj3H9C0
Nurses, cocoons, larvae and male:
https://www.youtube.com/watch?v=CfZq5iRGGKo
Individuals exploring the foraging area:
https://www.youtube.com/watch?v=vi1yFDwIbPI

#### Observational procedure of the static colony during resting periods

The experimental device for this experiment consists of two artificial nests (N1 and N2) of three rooms each (Room 1, 2 and 3) and a foraging area, linearly connected by six tunnels. At the beginning of the experiment, the brood is located in Nest 1. Each tunnel is equipped with two RFID readers that detect the passage and direction of tagged individuals between rooms.. The information recorded by readers are handled by two RFID electronics and then sent to a computer which creates and stores the data files.

The colony the 95 workers of the *Paraponera clavata* colony were monitored by RFID and by visual observation.

The colony was observed during two resting periods (first: from 13 January 13^th^ to January 19^th^, second: from January 25^th^ to February 9^th^).

30 Scan samplings were conducted on each one of these periods between 10h00 and 14h30.

We obtained by this way 60×95= 5700 individual observations conducing to a data set containing 5700 positions in the nest and 5700 behaviors visually observed.

In this paper we present the analysis of the observed behaviors in order to define the potential functional castes present in the colony. The data concerning the positions of the individuals in the nest obtained by observation and using the RFID apparatus are presented in the second part of this work in another publication.

For the data of the two resting periods, we conducted hierarchical ascendent classifications in order to obtain the clusters that could allow us to identify the different castes

Six categories of behaviors were observed, sampled and analyzed (Table 1).

**Table 1:** Behavioral categories observed during the resting periods

| Behavior | Description |
| --- | --- |
| Foraging | Explores or patrols in the nest or the foraging area |
| Eating | Eats in the foraging area where the meal dish were put |
| Collecting water | Takes water from the soaked papers available in the foraging area either to drink or to transport to the colony |
| Taking care of the brood | Touches an egg, a larva or a cocoon |
| Immobility | The ant does not move |
| Antenna contact | The ant touches its antenna with the antenna of another individual |

**Table 2:** Punctual transportation of material from the foraging area to the nest or to the garbage

| First resting period |  | First translocation |  | Second resting period |  | Second translocation |  |
| --- | --- | --- | --- | --- | --- | --- | --- |
| Number of transports | Number of individuals of each caste that did them | Number of transports | Number of individuals of each caste that did them | Number of transports | Number of individuals of each caste that did them | Number of transports | Number of individuals of each caste that did them |
| 51 | 4 nurses<br>16 intermed<br>4 foragers | 22 | 4 nurses<br>6 intermed<br>4 foragers | 34 | 2 nurses<br>3 intermed<br>9 foragers | 4 | 3 intermed<br>1 forager |

#### Observational procedure of the mobile colony during relocation experiments

The movement between nests of the colony of 95 individuals was observed during two emigration processes in experimentally induced relocations. Each worker had a colored tag attached to its thorax in order to be identify by the observer and by the RFID system

The first relocation was observed on January 20^th^ between 15h00 and 18h30

The second relocation was observed on February 9^th^ between 13h30 and 17h00

At _*time*_ = 0 we switch on a strong neon light over the first nest and we open the entrance of the second nest, then we record the colony movement until the entire brood is moved into the second nest.

During the relocation the observer notes the movement behaviors between the nests and for each individual she writes if the ant transports an egg, a larva, a cocoon or another ant from one room to another or from the original nest to the new nest.

The observer also writes if there are individuals who elicit olfactive marking behaviors (either putting olfactive traces in the ground with the extremity of the abdomen or spreading them with the abdomen).

#### Data analysis

For the two resting periods we conducted hierarchical ascendent classifications implemented in R in order to detect the potential clusters indicating the existence of functional castes using the data obtained by visual observation.

Concerning the data obtained by RFID the data files are in text format. They indicate, for each antenna scan (about three scans per second), the scan number, the date, time, and, for each individual (i.e., for each tag), which antenna is activated. If, during a scan, none is detected, nothing appears in the data file. If an ant moves from one room to another, it is detected by two successive antennas, and this allows us to infer the exact position of each ant at any moment.

## Results

The results concerning RFID monitoring were very similar to those observed by Cabanès and Benani (2010) for *Pachycondyla tarsata* indicating that the detection by the magnetic dispositive allows a very detailed monitoring of ants’s positions. As we mentioned before the results of the RIFD monitoring will be presented in another publication.

Interesting results were obtained by observing the particular behaviors of the ants

### Hierarchical ascendent analysis

The clustering analysis using hierarchical ascendant method allowed us to distinguish tree large clusters (Figure 5 and Figure 6).

**Figure 5:**
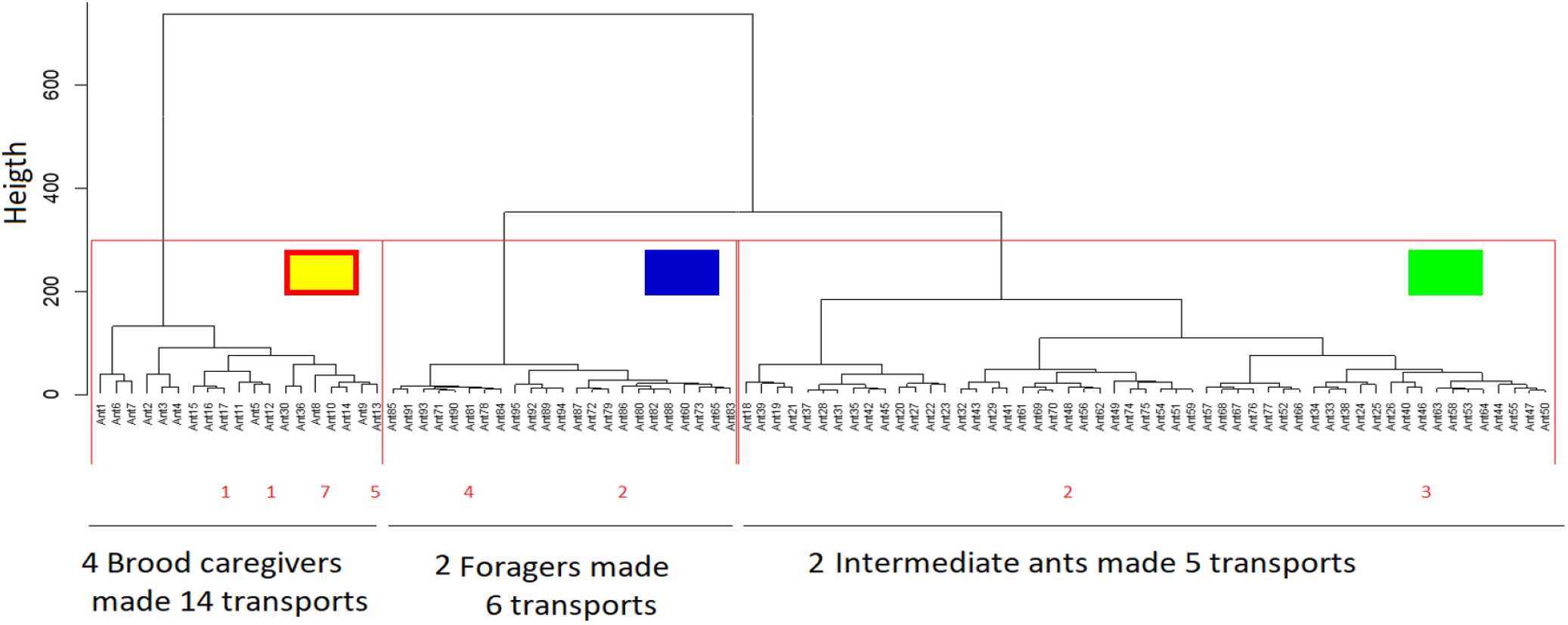
Clusters obtained by hierarchical classification of the behaviors observed in the 95 workers of the *Paraponera clavata* colony during the first resting period (red and yellow cluster: nurses, blue: predominant foragers, green: intermediate individuals)

**Figure 6:**
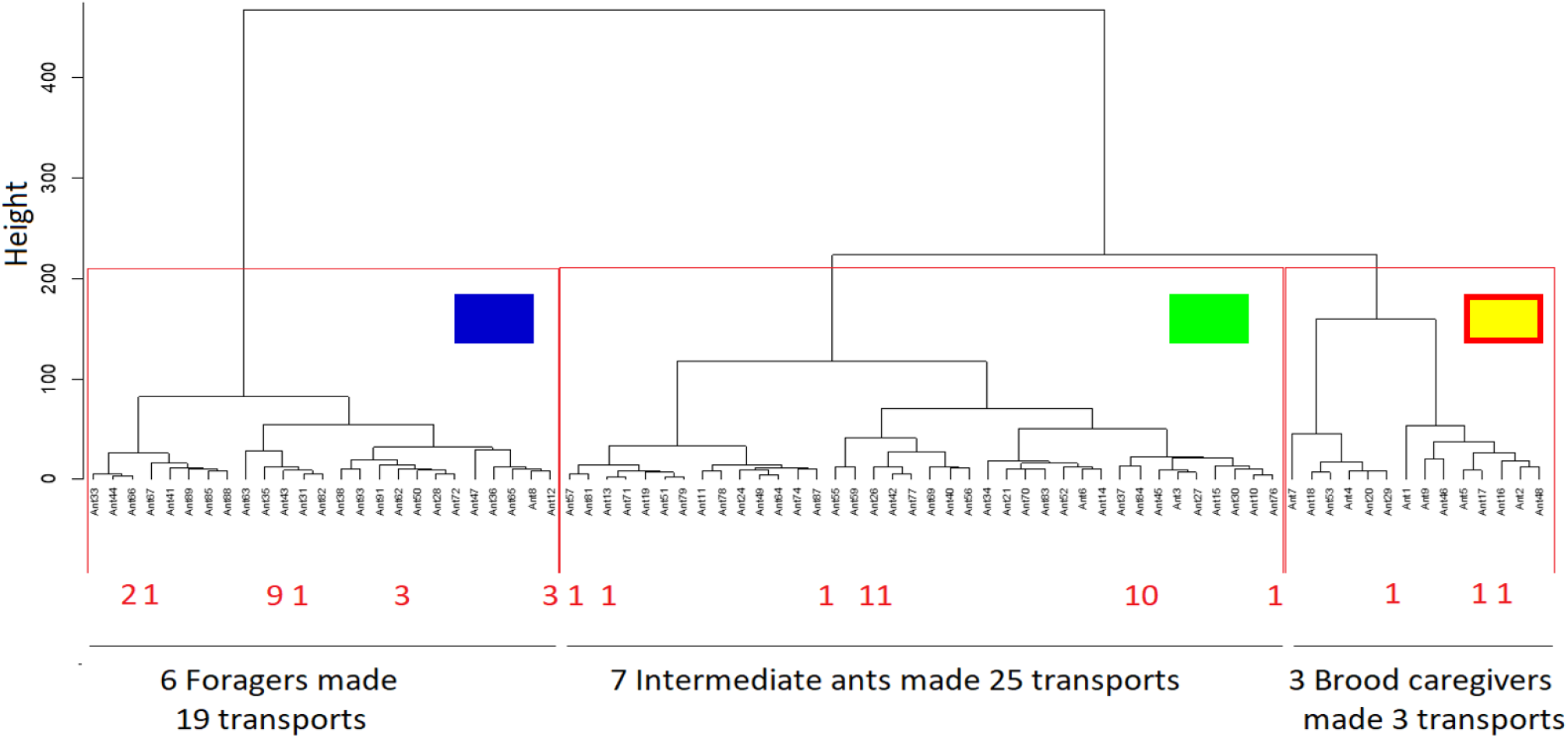
Clusters obtained by hierarchical classification of the behaviors observed in the 95 workers of the *Paraponera clavata* colony during the second resting period (red and yellow cluster: nurses, blue: predominant foragers, green: intermediate individuals)

The boxplots presenting the behaviors displayed by each cluster allow us to distinguish a first group of ants that significantly take care of the brood more often than the two other clusters (Figure 7) and who appeared to be very few times taking water from the paper probably because they spend most of their time in the nest and a restricted period in the foraging area. We can assume that these individuals represent the nurses caste of this colony during the observation period.

**Figure 7:**
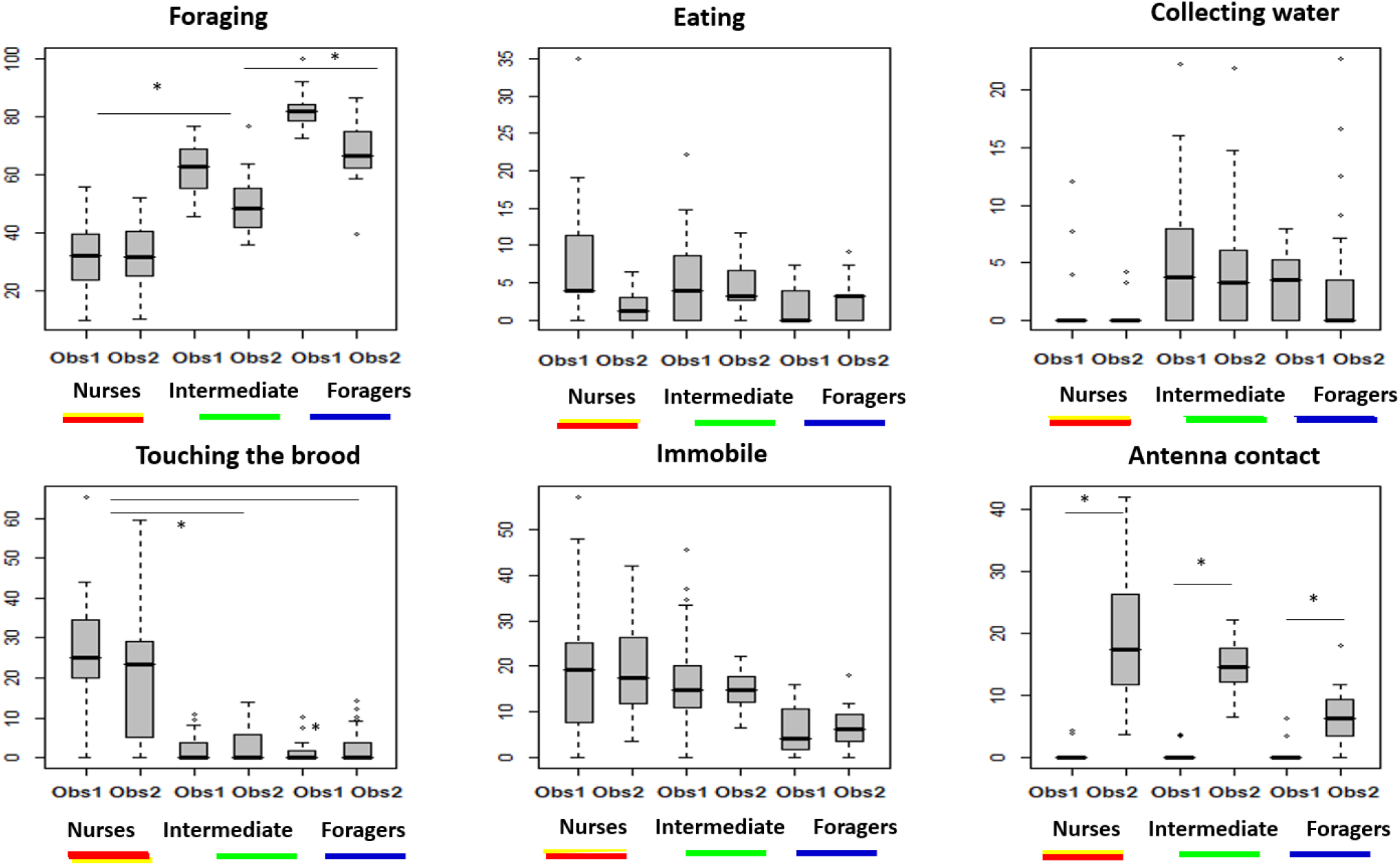
Behavioral profiles of the three clusters. Frequency of occurrence of six categories of behavior during the first resting period (Obs 1) and during the second resting period (Obs 2). Red and yellow: brood caregivers, Green: intermediate caste, Blue: foragers caste. (*: p<0.05 for Mann Whitney tests)

A second cluster (the blue one) was composed of individuals who forage significantly more than the individuals of the nurses’ cluster and than the individuals of the green cluster. By consequence we can assume that this group is the group of a specialized foraging caste that spent most of its time in explorative and patrolling behaviors. These observations are confirmed by the data indicating that they spend significantly less time immobile than the two other clusters. This group spent very little time taking care of the brood.

The third cluster (the green one) was composed of individuals that forage significantly more than the nurses but significantly less than the blue one. We can think that this is an intermediate caste that forages frequently but less than the clear foraging caste and that spent less time taking care of the brood than the confirmed nurses. Both the foraging cluster and the intermediate one was observed more often at the wet paper. This can be due to the fact that these individuals go to the paper in order to drink water but probably also to take water in their mandibles and transport it to the nest.

Concerning antenna contact for the three groups we observed that they displayed significantly more contacts of this type during the second resting period after the relocation. This is because the colony has to share many information concerning the new nest and the new situation of the colony or probably increases it grooming behaviors in order to socialize pheromones after this perturbation.

During the monitoring of the colony, we could observe some punctual behaviors as taking insects to the nest, transporting wet paper to the nest, transporting honey-apple, transporting death ants to the garbage, transporting waste to the garbage.

We present the distribution of these behaviors in the following table

9 different nurses from the 19 nurses transported material

24 different intermediate ants from the 53 intermediate ants transported material

17 different foragers from the 20 foragers transported material

Both in nurses and intermediate groups half of the group participates in the transportation of material. In the group of foragers almost al the individuals participate in the transports.

101 transports were observed. These behaviors were conducted by 8 ants that were nurses

In the following table (Table 3) we present the percentage of dead individuals between the first and the second resting period for each one of the castes.

**Table 3:**
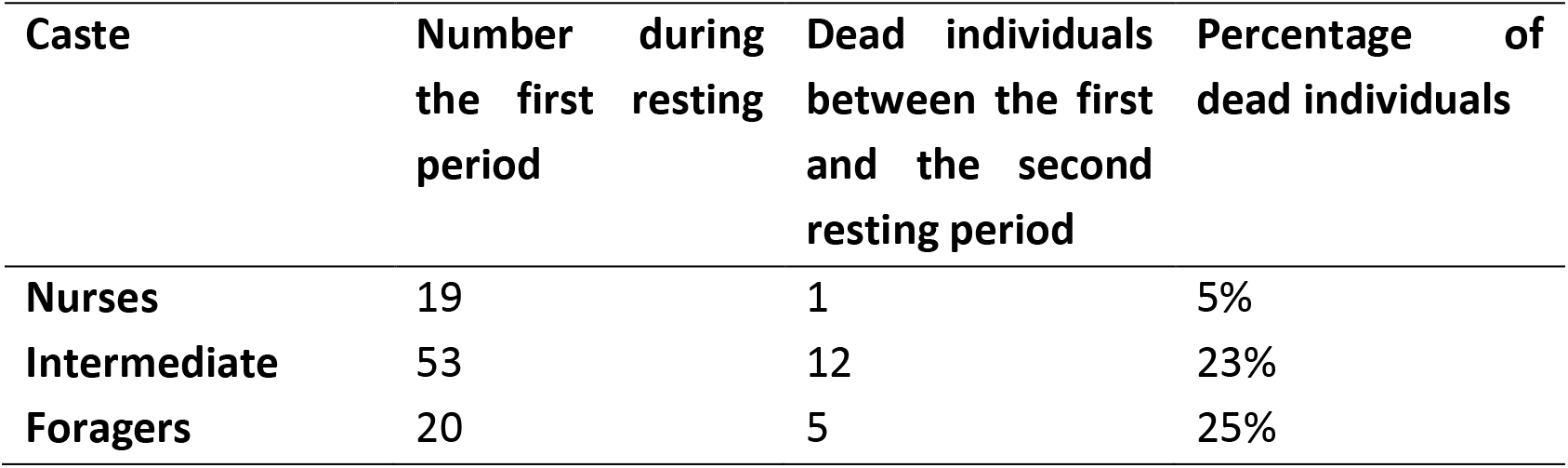
Dead individuals between the two resting periods

The proportion of dead individuals differed between the tree groups (χ^2^ test : p<0.05). taking into account the Bonferroni correction the limit for significancy here was 0.05/3= 0.017. The proportions of dead individuals were equivalent (χ^2^ test : p>0.05) between foragers and intermediate castes. The proportion of dead individuals was significantly higher in the intermediate caste than in the nurses’ caste p<0.017. The proportion of dead individuals was significantly higher in the foragers’ castes than in the nurses’ caste: p<0.017.

Stability of the roles and evolution in time

We compared the roles of the 76 ants that stayed alive between the first and the second resting periods and we observed that these roles were relatively stable.

44 of the 76 ants (58%) kept the same role (belonged to the same cluster) in the two resting periods.

29 of the 76 ants (38%) switched from one role to another role close to the first one (a forager became intermediate or vice-versa or a nurse became an intermediate or vice-versa)

3 of the 76 ants (4%) switched from nurse role to forager role.

7 nurses became intermediate ants, and 3 nurses became foragers between January 20^th^ and February 9th

12 intermediate ants became foragers between January 20^th^ and February 9^th^

These observations suggest that the most frequent evolution of ants’ status is:

nurse->intermediate->forager->death

However, we observed that 4 foragers became intermediate and 3 intermediate became nurses what seems to indicate the existence of a flexibility in roles.

Combining these information and data concerning deaths we can suggest that the organization of a *Paraponera clavata* colony evolves by this way (Figure 8):

**Figure 8:**
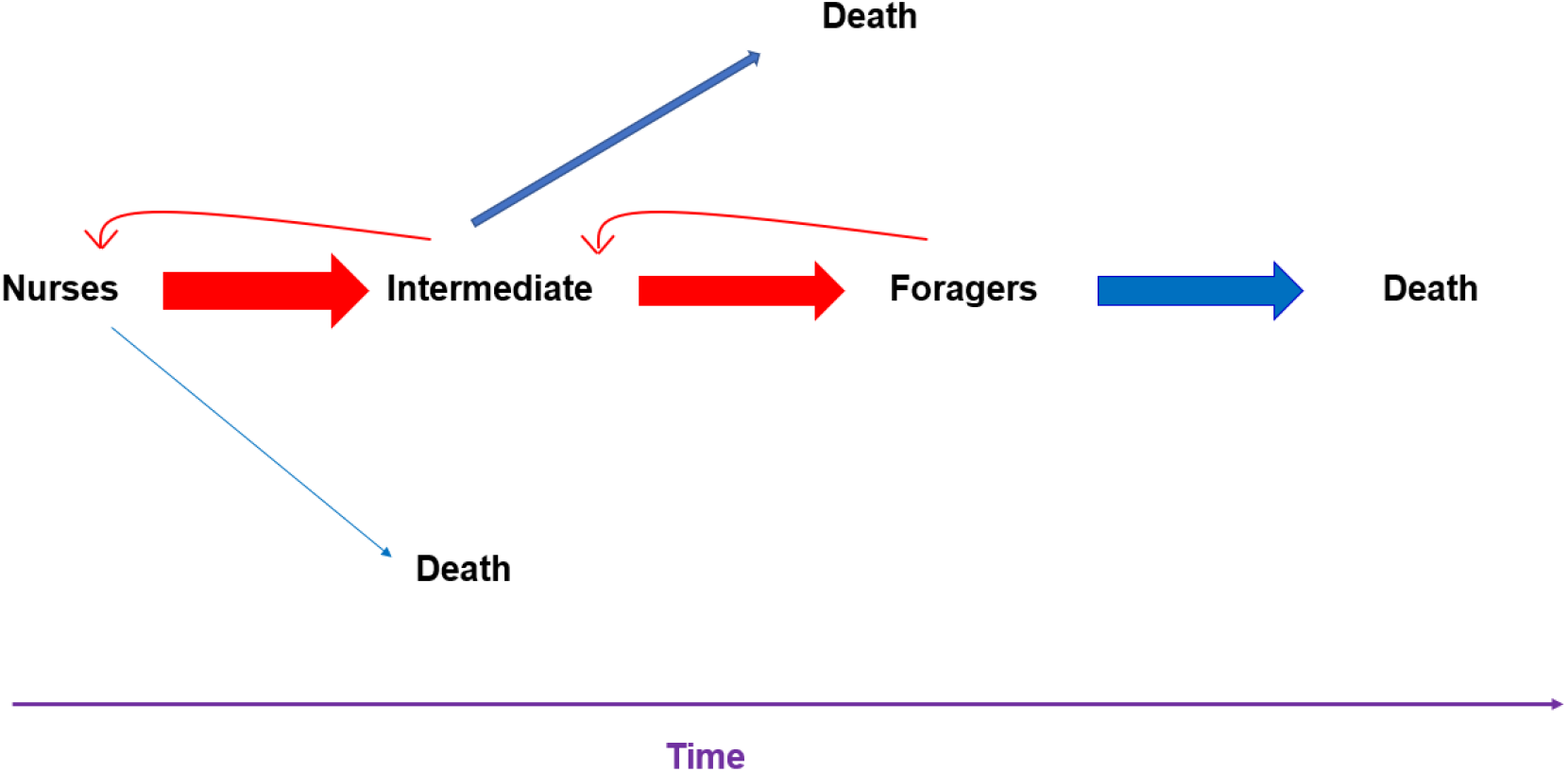
Evolution of the roles of the individuals in the colony

### Relocation

During the first relocation 25 ants were transported to the new nest by 8 of the 95 individuals of the colony (ie. 8.4% of the colony ensured the transport of the individuals that didn’t go to the new nest by their own). The transporters belonged to the tree castes. The number of individuals transported by each transporter ant was comprised between 1 and 7.

During the second relocation 47 ants were transported by 16 of the 78 individuals remaining in the colony (ie. 20.5% of the colony ensured the transport of the individuals that didn’t go to the new nest by their own). The transporters belonged to the tree castes. The number of individuals transported by each transporter ant was comprised between 1 and 11.

During the first relocation five larvae, 4 cocoons and one egg were transported to the new nest by 7 of the 95 individuals of the colony (ie. 7.3% of the colony ensured the transport of the brood. All the transporters belonged to the intermediate caste. The maximum number of items carried by each transporter was 3.

During the second relocation four larvae and three cocoons were transported to the new nest by 7 of the 78 individuals of the colony (ie. 9% of the colony ensured the transport of the brood. The transporters belonged to the three castes. The maximum number of items carried by each transporter was 3.

### Olfactive trails marking behaviors

The number of marking behaviors by individual were comprised between 0 and 30. 103 marking behaviors were observed during the first relocation.

5 of the 19 nurses elicit marking behaviors (44 in total). Thus 5 nurses (5.4% of the total number of individuals) made 43% of the olfactive traces. One of these nurses made 35 of the 44 marking behaviors.

11 of the 53 intermediate ants elicit marking behaviors (42 in total). Thus 11 intermediate ants (12% of the total number of individuals) made 41% of the olfactive traces.

11 of the 20 foragers elicit marking behaviors (17 in total). Thus 11 foragers made 16% (12% of the total number of individuals) of the olfactive traces.

Percentage of marking individuals in the nurse group: 26.3%

Percentage of marking individuals in the intermediate group: 20.7%

Percentage of marking individuals in the foragers group: 55%

The proportion of intermediate ants that mark is equivalent to the proportion of foragers that mark. However intermediate group marks two more times than foragers.

The nurse group marked as well as the intermediate group but 80% of the olfactive traces were put on the ground by one individual.

Nurses and intermediate ants ensured 84% of the chemical communication during the first relocation

Only 7 marking behaviors were observed during the second relocation

The colony marked 15 more times during the first than during the second relocation (X2: p<0.001) (Figure 9).

**Figure 9:**
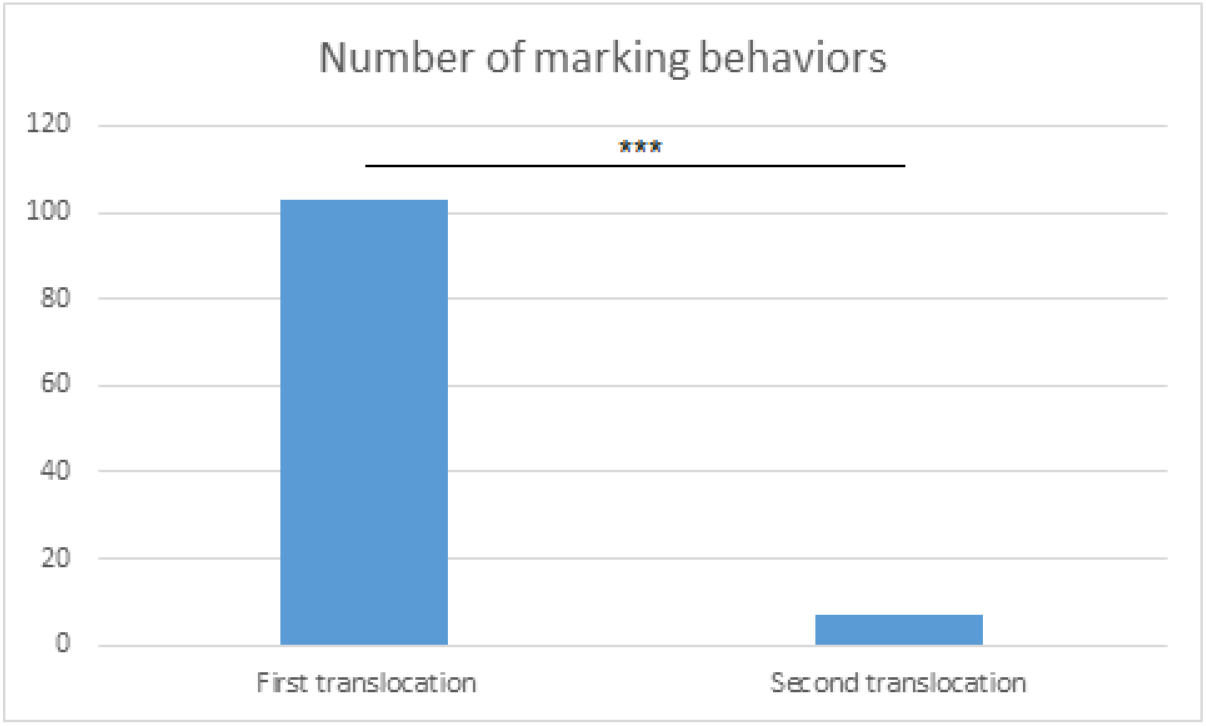
Number of marking behaviors during the first and the second relocation

## Discussion

From an ethological point of view, our results are of great help for understanding how tasks are distributed during resting periods and during nest relocation in *Paraponera clavata*. In fact, we obtained a very accurate description of the dynamic of the whole colony during all the two static periods and during emigration phases allowing us to emit some strong hypotheses about the function of the different individuals in the colonies of this ponerine species.

### Relevant results

1. The colony seems to be structured in three different functional castes: nurses (who take care of the brood and bring them water and food), intermediate ants and foragers (who forage, patrol and transport material between foraging areas and the nest)
2. The individuals of the colony seem to present an evolution in time from nurses to foragers with the intermediate status. This result is in agreement with the observations of other studies in ants where young individuals take care of the brood and older ones spend more and more time out of the nest foraging, providing resources to the the nest and patrolling until dead.
3. A flexibility in tasks is observed as some individuals can shift from intermediate status to nurse one or from forager status to intermediate one. This observation allows us to suggest that *Paraponera clavata* flexibility in tasks may confer some adaptability to the species and to allow adaptive reactions to environmental variability.
4. During the relocation processes the brood is carried to the new nest and put in safe by individuals belonging to the tree castes indicating that the priority in critical situations is to relocate the vulnerable brood rapidly as there is no evidence of caste specialization for this task.
5. During the relocation food and water is transported to the new nest by the tree castes suggesting that the reorganization of a suitable nest is a priority to which all the castes contribute.
6. Marking behaviors were 15 times more frequent in the first relocation (from nest 1 to nest 2) than in the second relocation (from nest 2 to nest 1). This can be because the first relocation is a really novel situation for this colony and much information to join the new nest is needed. On the contrary, during the second relocation the colony moves to a nest that has been already occupied before and for which the travel is known.
7. Intermediate individuals work double than foragers during olfactive deposition indicating that this caste is probably relevant for communication signaling during relocation events.
8. During the first relocation the transportation of most of the individuals that did not walk and that were carried to the new nest was ensured by the nurses and the intermediate ants. In the second relocation these transports were conducted by intermediate and forager ants. We can suggest that intermediate ants play an important role in this aspect and that nurses and foragers alternatively transport the individuals that do not walk as a way to share the labor from one relocation from another. The individuals that do not walk may be young individuals that have no experience in going out of the nest or individuals scared by the new situation. In both relocations the male didn’t want to leave the nest. He was grabbed to the floor and another ant must remove him from the old nest and transport him to the new one.
9. It seems that there is no caste specialization in ants transport during relocations, but some particular individuals made many of these transports probably the stronger or the most experienced ones.
10. We detected different functional roles in this species who does not have clear morphological castes. This finding is relevant as it let us suggest that division of labor is not only present in species that have morphological polymorphism but also in species were individuals are apparently homogenous.

Some results are in accordance with previous works, especially the behaviors that can be observed in the foraging area. For example, the dynamic of the transportation behavior match up the results presented in Pezon et al. (2005). These hypotheses should now be validated by repeating the experiment with different colonies and different species. A complete understanding of the emigration process based on systematic experimentations would be an important step ahead for the research in *Paraponera clavata* and other social insects. Individuals of this species build underground nests with intercommunicated chambers on surfaces as big as 1200m3, in a 130m tunnel between several chambers has even been observed (Braun et al., 1994). Peripheral nests constitute advanced positions which strengthen the network. The queen and brood are particularly mobile in order to optimize brood rearing. In the wild, solitary foragers look for termites through visual orientation (Hölldobler 1980). They also exhibit mass recruitment through trail pheromones when an important food source is discovered (Hölldobler 1984) and prey transport is allocated between different foragers who do only part of the return trip. Workers also collect insect remains and contribute to resource turnover (Dejean et al. 1993). It is therefore an interesting model for laboratory studies of colony movements.

Our laboratory results are in agreement with these field observations and suggest that even if there is a castes organization structure in this species the flexibility and alternative repartition of labor may facilitate the functioning of the colony as efforts and energetic investment are shared.

#### Perspectives

During our experiments we ensured an RFID monitoring which conduced to the registration of a big data set concerning all the ants’ positions during the observation phases. In a further publication we will present the detailed positional dynamics and movements of the ants during resting and relocation periods. The abstracted individual activity during emigration can be used to compute the collective dynamic of emigration. We think that these dynamics would be similar to those observed in a previous study on *Pachycondyla tarsata* (Cabanès et al. 2009) as presented below (Figure from Galassi et al. 2009)

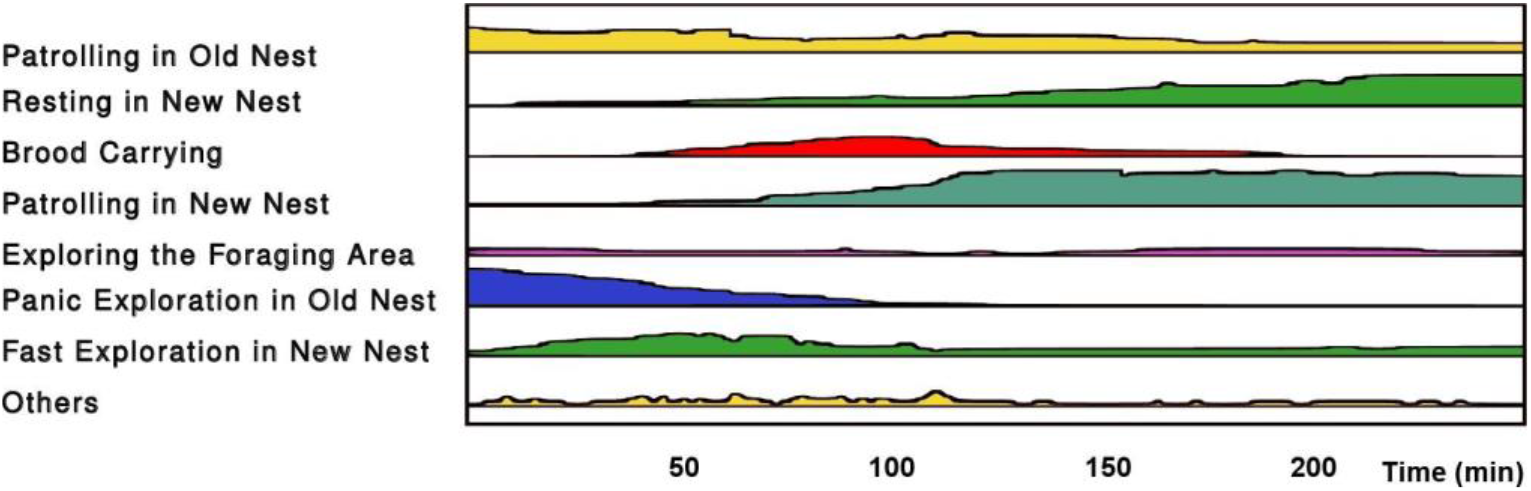

Activity analysis: evolution of the number *n_ij_* of individuals involved in the activities *A*_0_…*A*_2_.

This figure is a representation of this dynamic, showing the number of ants expressing each activity during the migration process. As we can see, colony emigration follows a typical pattern: when the light is switched on, the first event is a panic exploration of the old nest that is expressed by most of the ants. Slower patrolling will remain constant during all the process and concern not only the old nest but also the foraging area. The second event is the discovery of new nests followed by the brood transport. Afterwards, a more constant exploration of the new nest occurs. The last activity, which appears gradually, is the settlement in the new nest.

The analysis of the recordings made on *Paraponera clavata* with RFID methods and more studies on these phenomena would allow the myrmecologist community to better understand the structures and dynamics of these amazing eusocial evolved group.

